# Proteomic analysis of coarse and fine skin tissues of Liaoning cashmere goat

**DOI:** 10.1101/2021.09.20.461155

**Authors:** Zhixian Bai, Yanan Xu, Ming Gu, Weidong Cai, Yu Zhang, Yuting Qin, Rui Chen, Yinggang Sun, Yanzhi Wu, Zeying Wang

## Abstract

Proteomics is the study of all proteins expressed by a cell or even an organism. However, knowledge of proteins that regulate the fineness of cashmere is limited. Liaoning Cashmere goat (LCG) is a valuable genetic resource of China. The skin samples of Liaoning cashmere goats during the growing period were collected performed Tandem Mass Tag (TMT) method and identified 117 differentially expressed proteins in CT_LCG (course type) and FT_LCG (fine type). To verify protein genes differentially expressed in LCG, we performed PRM validation on three candidate proteins (*ALB*, *SDC1* and *ITGB4*) in CT-LCG and FT-LCG. Furthermore, primary metabolic process and lysosome are most enriched in the GO and KEGG pathways, respectively. In addition, we also derived a protein-protein interaction (PPI) regulatory network from the perspective of bioinformatics. This study sought to elucidate the molecular mechanism of differential proteins regulating cashmere fineness of Liaoning cashmere goats by using TMT quantitative proteomics analysis. Differentially expressed proteins *ALB* and SDC1 may regulate cashmere fineness, *ITGB4* can be further studied as a promising protein. They can be used as key genes to lay a foundation for the study of cashmere fineness of Liaoning cashmere goats.

## Introduction

Cashmere goats are primarily bred for cashmere woold production[1,2], and it is one of the most important genetic resources in China[3]. Cashmere is an important textile with high economic value and is known as “soft gold” in the world[4,5]. With the development of cashmere processing industry, the cashmere market is in short supply and the price is rising. In the face of the increasing consumption demand at home and abroad, the production and scientific research of cashmere in China have also achieved beneficial development[6]. The fineness and yield of cashmere are important factors to enhance the economic value of cashmere. With the increasing demand of cashmere goats for cashmere year by year, the cultivation of superfine quality cashmere goats has become a pressing problem to be solved in cashmere goat breeding[7].

Proteomics is essentially the large-scale study of the properties of proteins, proteins are the basis for animal-derived filaments and hair fibers[8]. The importance of proteomics in animal science and its applications has increased significantly in recent years, but there remain certain limitations, particularly in terms of public databases[9]. In this regard, proteomics-derived data is important in sequencing and further study of species, including sheep[10]. Mammalian hair fibers are complex and include a large number of keratin and keratin-associated proteins [11], as well as lipids and carbohydrates [12]. Several proteins have been found in studies that contribute to the composition and structure of wool [13,14] and can also be found in dissected hair follicles [15]. Proteomics studies include dynamic description of gene regulation, the quantitative determination of gene expression protein levels, the identification of diseases, drugs on the impact of life processes, and the interpretation of gene expression regulation mechanisms. Therefore, we speculate that proteomics plays a key role in the fineness of cashmere wool. Parallel reaction monitoring (PRM) is a kind of ion monitoring technology (5-6 orders of magnitude), it’s in high precision parallel testing all product ion mass spectrometer[16]. Through the PRM to validate the low abundance of the differentially expressed proteins, research on the effect of the cashmere protein mechanism, improve cashmere fineness.

To further understand the proteins regulating cashmere fineness, the mechanism of cashmere fineness regulation was studied in this study. Our aim was to investigate the function of proteins and related genes in LCG. We quantitated the skin differentially expressed proteins of LCG using PRM, and identified key candidate proteins related to cashwool fineness by gene ontology (GO) and Kyoto Encyclopedia of Genes and Genomes (KEGG) enrichment analyses. The molecular regulation mechanism of cashmere fiber growth has not been fully established. This study provides a helpful reference for further understanding the association between cashmere fineness and proteomics, and also provides a reference for understanding the development of cashmere character.

## Materials and Methods

### Animal Ethics

Animal in the whole process of experiments were on the basis of Animal Experimental Committee of Shenyang Agricultural University (Shenyang, China, 201906099). In our study, these activities do not require specific permits and the animals do not involve endangered or protected species.

### Experimental animals and management

Sheep shed high and dry terrain, to the sun and leeward, sheep shed before leaving 10-20 square meters of feeding area, installed fixed feeding trough and drinking water appliances. Sheep house temperature 4-7 °. Each sheep has an area of 1.5 to 2.5 square meters. Sheep are mainly to eat coarse feed, concentrate as a supplement. The selected sheep are healthy and free from disease infection.

### Total Protein Extraction and Quality Test

The samples were ground in liquid nitrogen, cracked with buffer, and then ultrasonic on ice for 5 min. The lysate was centrifuged at 12000 g at 4° for 15 min, and the supernatant was transferred to a clean tube. The extracts of each sample were reduced at 56° C for 1h with 10 mM DTT and then alkylated at room temperature in the dark for 1h with sufficient iodoacetamide. The samples were then vortically mixed with 4x volume of precooled acetone, incubated at −20° for at least 2 h, and centrifugated to collect the precipitation. After two washes with cold acetone, particles were dissolved in a solution buffer consisting of 0.1m trisethylammonium bicarbonate (TEAB, pH 8.5) and 6m urea.

Using Bradford Protein quantification kit, prepare BSA standard protein solution according to the instruction, concentration gradient range is 0-0.5 *μ* g/*μ* L. The BSA standard protein solutions with different concentration gradients and the sample solutions with different dilution ratios were added to the 96-well plate, and the volume was supplemented to 20 *μ* L, and each gradient was repeated 3 times. 180 *μ* L G250 solution was quickly added and placed at room temperature for 5 min. The absorbance at 595 nm was determined. The absorbance of standard protein solution was used to draw the standard curve and calculate the protein concentration of the sample to be tested. 12% SDS-PAGE gel electrophoresis was performed on 20 *μ* g protein samples of each sample. The gel electrophoresis strips were 80 V and 20 min, and the gel electrophoresis conditions were 120 V and 90 min. After electrophoresis, coomassie bright blue R-250 staining was performed until the bands were clear.

### TMT Labeling of Peptides

Take 120 *μ* g of each protein sample, add 1 *μ* g/*μ* L trypsin 3 *μ* L and 50 mM TEAB buffer 500 *μ* L, mix and digest overnight at 37°. An equal volume of 1% formic acid was mixed with the digested sample and centrifuged for 5min at 12000 g at room temperature. The supernatant was slowly supernatated onto the C18 column, washed with 1mL washing solution (0.1% formic acid, 4% acetonitrile) for 3 times, and then eluted with 0.4ml eluting buffer solution (0.1% formic acid, 75% acetonitrile) for 2 times. The eluent of each sample was combined and lyophilized. Add 0.1 M TEAB buffer 100 *μ* L to recombine, add TMT labeling reagent 41 *μ* L dissolved in acetonitrile, shake the mixture at room temperature for 2 h. The reaction is then terminated by adding 8% ammonia. All labeled samples were equally mixed, desalted and lyophilized.

### LC-MS/MS Analysis

To construct the transition library, shotgun proteomics analysis was performed using the Easy-NLCTM 1200 UHPLC system (Thermo Fisher) and QEXactive HF mass spectrometry (Thermo Fisher) operating in data Dependence Acquisition (DDA) mode. One sample was injected into a self-made C18 Nano-Trap column (2cm×75 *μ* m, 3 *μ* m). In a self-made analytical column (15 cm× 150 *μ* m, 1.9 *μ* m), linear gradient elution was used. The separated polypeptides were analyzed by q-Exactive HF mass spectrometer (Thermo Fisher). The ion source was Nanospray FlexTM(ESI), the spray voltage was 2.3 kV, and the ion transport capillary temperature was 320°C. The full scan range is from M /z 350 to 1500, with a resolution of 60000 (m/z 200), automatic gain control (AGC) target value of 3 × 106, and a maximum ion implantation time of 20 ms. The top 20 predecessors have the highest rich selection of full scan and dispersed energy collision dissociation (HCD) and MS/MS analysis, resolution 15000 (M/Z 200) 6-plexus (45,000 10-plexus, M/Z) 200), the target value of automatic gain control (AGC) is 5× 104, the maximum ion implantation time is 45 ms, the normalized impact energy is 32%, the intensity threshold is 1.9 × 105, and the dynamic exclusion parameter is 20 s.

### Data analysis

The spectra of each fraction were searched separately with NCBI database by the search engine Proteome Discoverer 2.2 (PD 2.2, Thermo). The protein contains at least one unique peptide with FDR not exceeding 1.0%. Proteins containing similar peptides that could not be distinguished by MS/MS analysis were identified as the same proteome. Reporter quantification was used for TMT quantification (TMT 10-plex). Protein quantitative statistical analysis of testing results the Mann - Whitney, quantitative results in a significant difference between experimental group and the control group (p < 0.05 and | log2FC | > 1.2 or < 0.83) of the protein is defined as the differentially expressed proteins (DEP).

Non-redundant protein databases (including Pfam, PRINTS, ProDom, SMART, ProSiteProfiles, PANTHER)[40] and Clusters of Orthologous COG were added by interProSCAN-5 program Groups) and KEGG (Kyoto Encyclopedia of Genes and Genomes) databases for analyzing protein families and pathways. Possible protein-protein interactions were predicted using the String-DB server [41](http://string.embl.de/). Enrichment analysis of go, IPR and KEGG was carried out using [42] enrichment pipeline.

### PRM Validation

To verify the protein expression level obtained by TMT analysis, the selected proteins were analyzed by LC-PRM MS in Shanghai Applied Protein Technology Co., LTD[43] to further quantify the expression level. Peptides were prepared according to TMT protocol and AQUA stable isotope polypeptides were added to each sample as internal standard reference.Tryptic peptide was loaded onto the C18 stage tip for desalination prior to reversed-phase chromatography on the Easy NLC-1200 system (Thermo Scientific). Gradient liquid chromatography with acetonitrile for 1 h, gradient range of 5-35%, 45 min. Q Exactive Plus mass spectrometer (Thermo Scientific) was used for PRM analysis. In the experiment, a unique peptide with high strength and high confidence for each target protein was used to optimize the impact energy, charge state and retention time of the peptide that most significantly regulated it. The mass spectrometer operated in positive ion mode with the following parameters: MS1 full scan was obtained with a resolution of 70,000 (200 m/z), automatic gain control (ACG) target value of 3.0×10-6, and maximum ion implantation times of 250 ms. The full MS scan was followed by 20 PRM scans with 35000 resolution (m/z 200), AGC 3.0×10-6, and maximum injection time 200 MS. The target peptide was isolated through the second window. In high Energy dissociation (HCD) collision cells, ion activation/dissociation takes place at a standardized collision energy of 27. The raw data were analyzed using Skyline (MacCoss Lab, University of Washington)[44], where the signal strength of the individual peptide sequence of each significantly altered protein was quantified relative to each sample and normalized to a standard reference.

## Result

### Identificaiton of proteins in CT_LCG and FT_LCG

TMT analysis in this study showed that 70,066 spectrograms were matched out of 314,881 secondary spectra. A total of 29446 peptides and 4022 specific proteins were identified, and 3999 proteins were quantified from all samples(Table 1). The molecular weight of these proteins is mainly above 100 kDa(figure 1A). The detection of more peptides associated with a given protein, the more reliable its identification. As such, the coverage of protein identification can provide insight into overall identification result accuracy. The results showed the identification of 79.61% of the proteins, with >10% peptide sequence coverage. Identification was achieved for 65.49% of the proteins, with >20% peptide sequence coverage (figure 1B). Therefore, the protein identification in this study was accurate, which provided a basis for the selection of differential proteins.

**Table 1.** Protein identification

| Run name | Total spectra | Matched spectrum | Peptide | Identified protein | ALL |
| --- | --- | --- | --- | --- | --- |
| run | 314881 | 70066 | 29446 | 4022 | 3999 |

**Figure 1.**
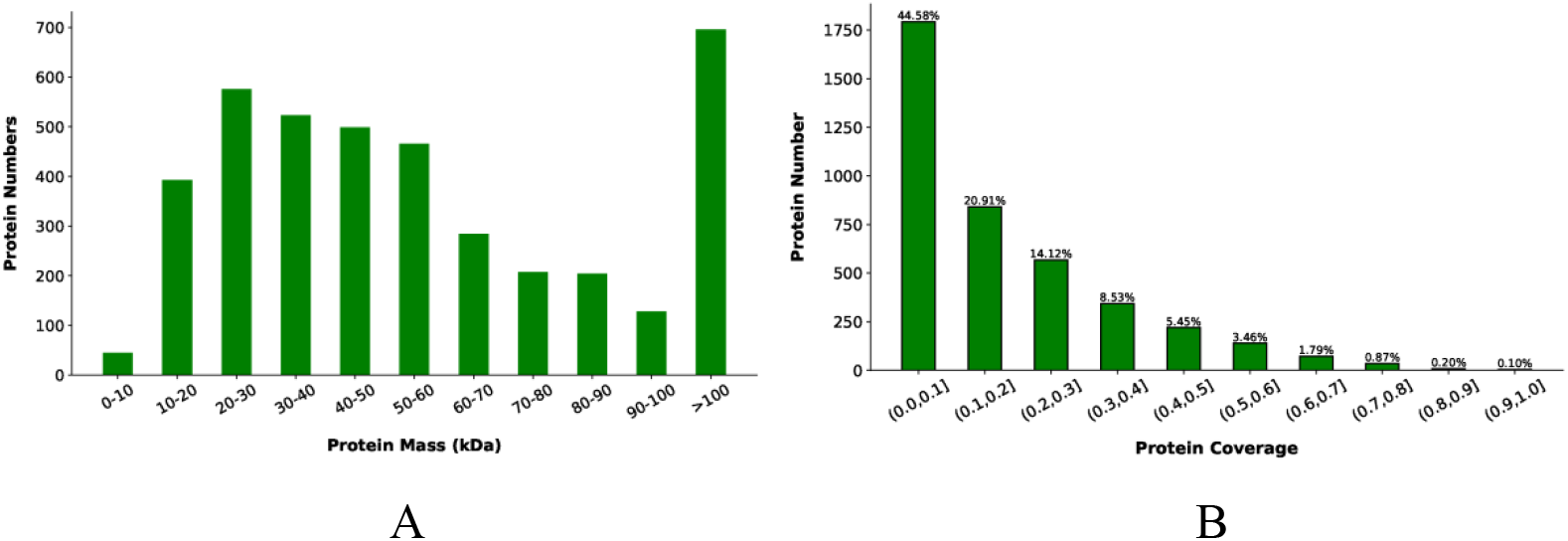
Identificaiton of proteins in CT-LCG and FT-LCG. (A)Molecular weight distribution of protein.The abscissa is the molecular weight of the identified protein, and the ordinate is the quantity of the identified protein. (B) Distribution of Protein Coverage. The abscissa is the interval of protein coverage: the length of the protein covered by the detected peptide/the total length of the protein, and the ordinate is the number of proteins contained in the corresponding interval.

### Total protein functional classification and pathway analyses

According to GO, proteins are divided into three regions :856 biological processes, 834 cellular components and 1,962 molecular functions(figure 2A). Cellular component terms were most enriched. To examine the efficacy of our labeling strategy, the COG database was leveraged to classify putative protein functions. The most common categories were General function prediction only; Posttranslational modification, protein turnover, chaperones; and Translation, ribosomal structure and biogenesis (Figure 2B). KEGG pathway analyses for these proteins were also performed (Figure 2C). Interestingly, many proteins are annotated in the global and overview diagrams.

**Figure 2.**
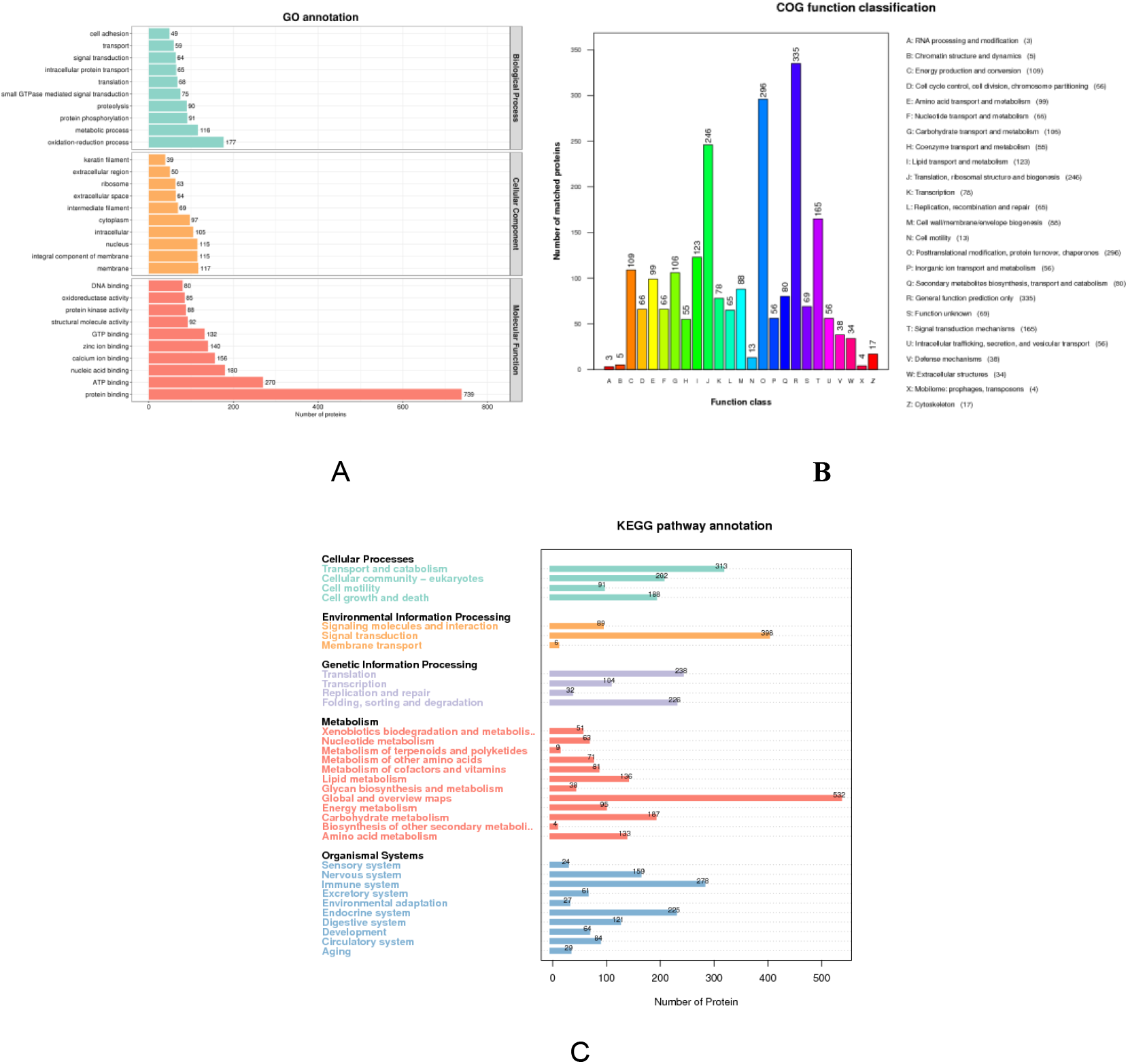
Function distribution of total proteins. (A) GO term classifications. (B) COG functional classification. (C) KEGG pathway annotations.

**Fig.2.**
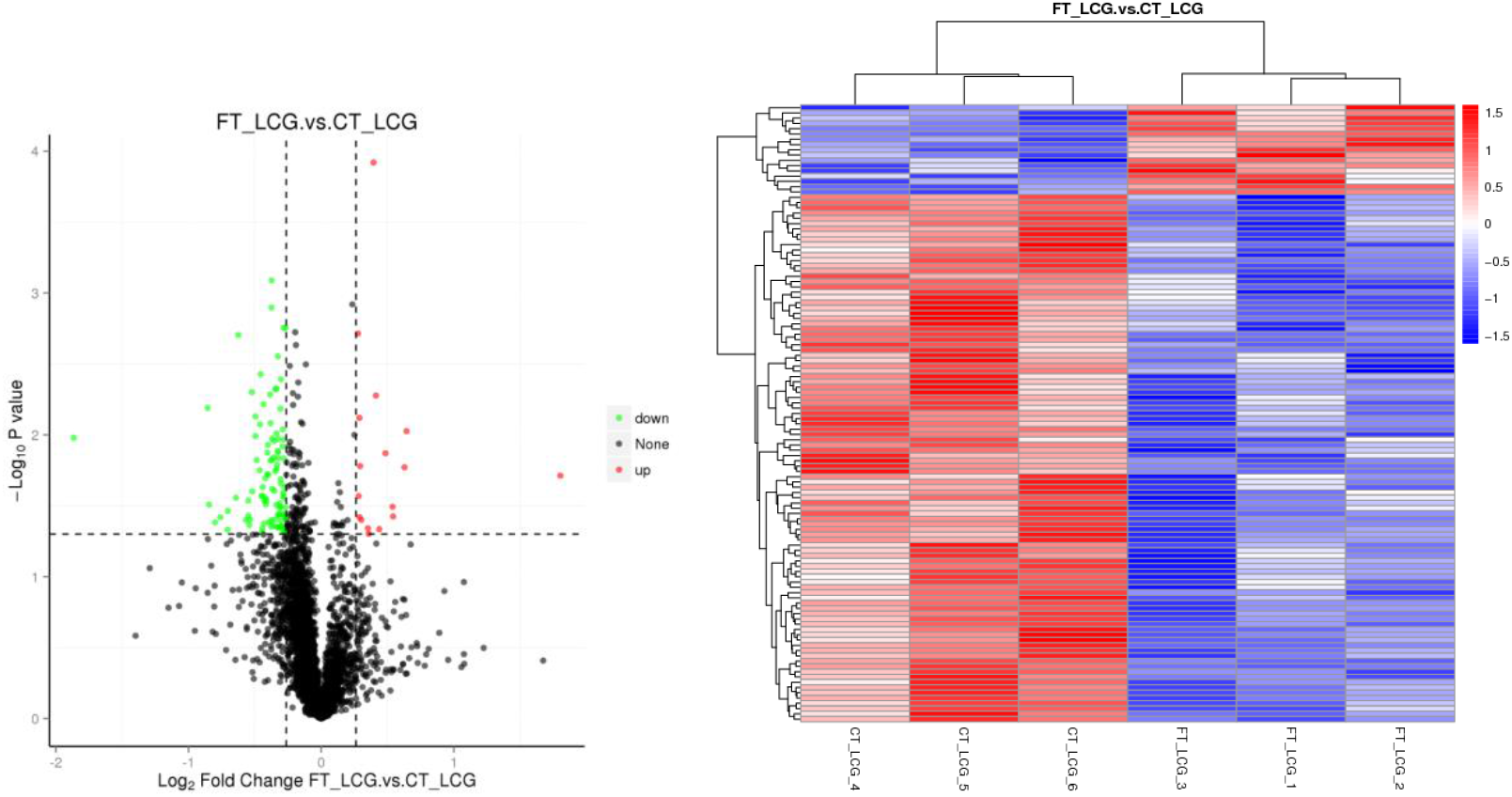
(A)Volcano map of differentially expressed proteins. The abscissa represents the difference multiple of the differential protein (log2 value), and the vertical axis represents Pvalue (−log10 value). Black represents the protein with no significant difference, red represents the up-regulated protein (FC≥1.2,Pvalue≤0.05), and green represents the down-regulated protein (FC≤0.83,Pvalue≤0.05). (B)Heat map of differentially expressed proteins. Rows correspond to individual proteins, with red and blue respectively indicating higher and lower relative abundance.

### Identificaton of differential proteins in CT-LCG and FT-LCG

We identified 117 expressed proteins that different significantly between coarse and fine types of Liaoning cashmere goats. Of these, 17 and 100 proteins were up- and down-regulated, respectively, in coarse types of Liaoning cashmere goats compared with fine types. Top20 was selected, as shown in Table 2. Volcano plots representing these proteins were prepared (Figure 2A), with differentially expressed proteins (≥1.2-fold, P<0.05) being located in the upper quadrant. Hierarchical clustering analysis was performed for the differentially expressed proteins to better display protein abundance differences between groups(figure 2B).

**Table 2.** Differentially expressed proteins with target genes in CT-LCG and FT-LCG

| Protein | Gene | FT_LCG<br>average | CT_LCG<br>average | FT_LCG.vs.CT_LC<br>G FC | FT_LCG.vs.CT_LC<br>G P-value | FT_LCG.vs.CT_LC<br>G log2FC | FT_LCG.vs.CT_LC<br>G UP.DOWN |
| --- | --- | --- | --- | --- | --- | --- | --- |
| NP_851335.1 | ALB | 205.4 | 109.6 | 1.45592852 | 0.037605649 | 0.541939527 | up |
| XP_017911091.1 | SDC1 | 107 | 162.3 | 0.767685403 | 0.015152173 | -0.381412879 | down |
| XP_024852938.1 | COL6A5 | 177.1 | 267 | 0.774143302 | 0.024054013 | -0.369327446 | down |
| XP_024835563.1 | ITGB4 | 423.3 | 562.6 | 0.822911099 | 0.027897607 | -0.281191513 | down |
| XP_017895824.1 | GUSB | 4007 | 8558 | 0.61324073 | 0.046601379 | -0.705474574 | down |
| XP_005227352.1 | CTSD | 585.4 | 1818.8 | 0.274607424 | 0.010483377 | -1.864557465 | down |
| XP_017907267.1 | CTSB | 5566.1 | 5956.5 | 0.805688817 | 0.035963673 | -0.311705364 | down |
| XP_017919849.1 | BLMH | 7015.8 | 7982.4 | 0.78158 | 0.043533849 | -0.355534544 | down |
| XP_005689186.1 | FABP9 | 6717.8 | 9203.4 | 0.709529151 | 0.007418649 | -0.495066135 | down |
| XP_005692366.2 | DMKN | 4209.4 | 6582.9 | 0.726373504 | 0.008433917 | -0.461216517 | down |
| XP_017916681.1 | LOC106503064 | 32919.9 | 17336.4 | 1.545767927 | 0.01693558 | 0.628323737 | up |
| XP_017913428.1 | CTSA | 4516 | 6079 | 0.728864107 | 0.003733355 | -0.456278238 | down |
| XP_005683176.2 | LOC102174022 | 3141.7 | 4619.8 | 0.765517347 | 0.005200705 | -0.385493025 | down |
| NP_001094595.1 | ENO2 | 179.3 | 197.3 | 0.783634933 | 0.024820504 | -0.351746383 | down |
| XP_005204963.1 | LOC510369 | 233.8 | 298.7 | 0.749069818 | 0.030975627 | -0.416827901 | down |
| XP_017909238.1 | REEP5 | 5968.8 | 5276 | 1.212198862 | 0.001930232 | 0.277626394 | up |
| XP_005700873.1 | RGN | 1426 | 2153.5 | 0.674784217 | 0.039837184 | -0.567501865 | down |
| XP_017903944.1 | PLBD1 | 3167.1 | 4371.7 | 0.709320118 | 0.010186335 | -0.495491228 | down |
| XP_017901365.1 | LOC102185621 | 2066.8 | 2714.9 | 0.687001385 | 0.043275425 | -0.541615088 | down |
| XP_005691900. | CRISPLD2 | 1500.7 | 1838.5 | 0.735027667 | 0.048232908 | -0.44412954 | down |

### Differentially expressed protein functional classification and pathway analyses

Differentially expressed proteins were next subjected to GO term enrichment analyses as above. Biological process analyses revealed these proteins to be primarily associated with the primary metabolic process. Cell component analysis showed that these proteins were primarily localized in the plasma membrane and the whole plasma membrane. In addition, molecular functional analysis showed that these proteins were mostly associated with hydrolase activity(Figure 3A). The most significantly enriched KEGG pathway was lysosome (Figure 3B). The subcellular localization ratio of different proteins in each comparison was statistically analyzed, and the proportion of nuclear protein, cytoplasmic protein and extracellular protein accounted for more than 50%.

**Fig.3.**
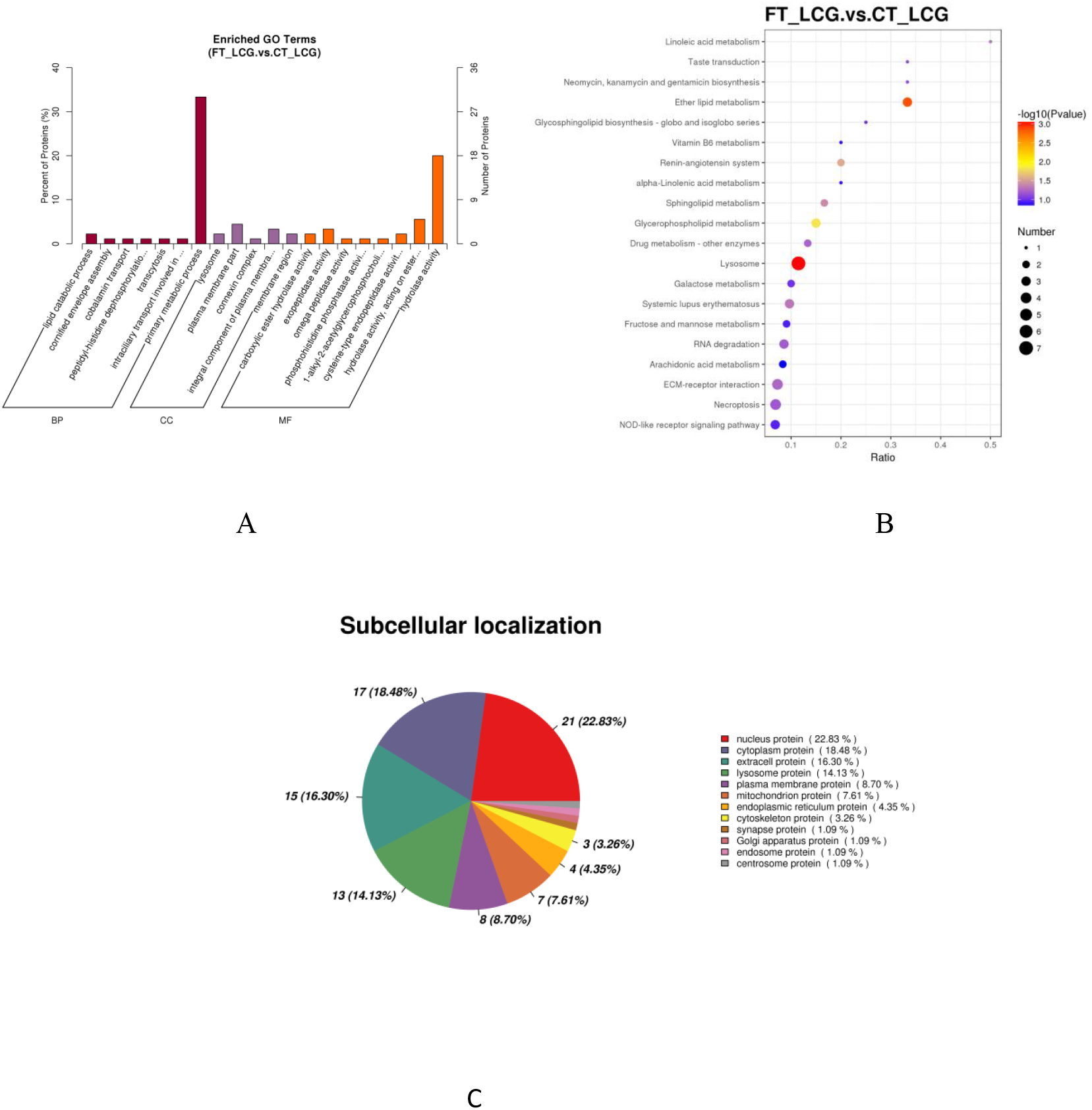
Functional Analyses. (A) Go enrichment. The figure shows the enrichment results in three categories, each with a maximum of 20 (Pvalue≤0.05). The percentage of vertical axis represents x/n in the table. (B) KEGG enrichment. The abscissa in the figure shows the ratio of the number of differential proteins in the indicated pathway relative to the total protein number associated with that pathway. Greater values indicate a higher degree of differential protein enrichment in the pathway. (C) Subcellular localization analysis.The number and distribution ratio of proteins in each suborganelle are shown in a pie chart.

### Validation of Proteins by PRM

During validation, we assessed the number of 9 candidate proteins and determined the expression diversity among CT_LCG ang FT_LCG samples. Three targeted proteins in LCG were quantified by PRM (Figure 4). The quantitative validation of PRM method has good correlation with TMT results. The findings showed that ITGB4 promising protein showed similar up-regulation or down-regulation in both TMT and PRM pathways. Interestingly, ALB was significantly upregulated and SDC1 was significantly downregulated in FT_LCG compared to CT_LCG, whether using TMT or PRM(table 3).

**Fig.4.**
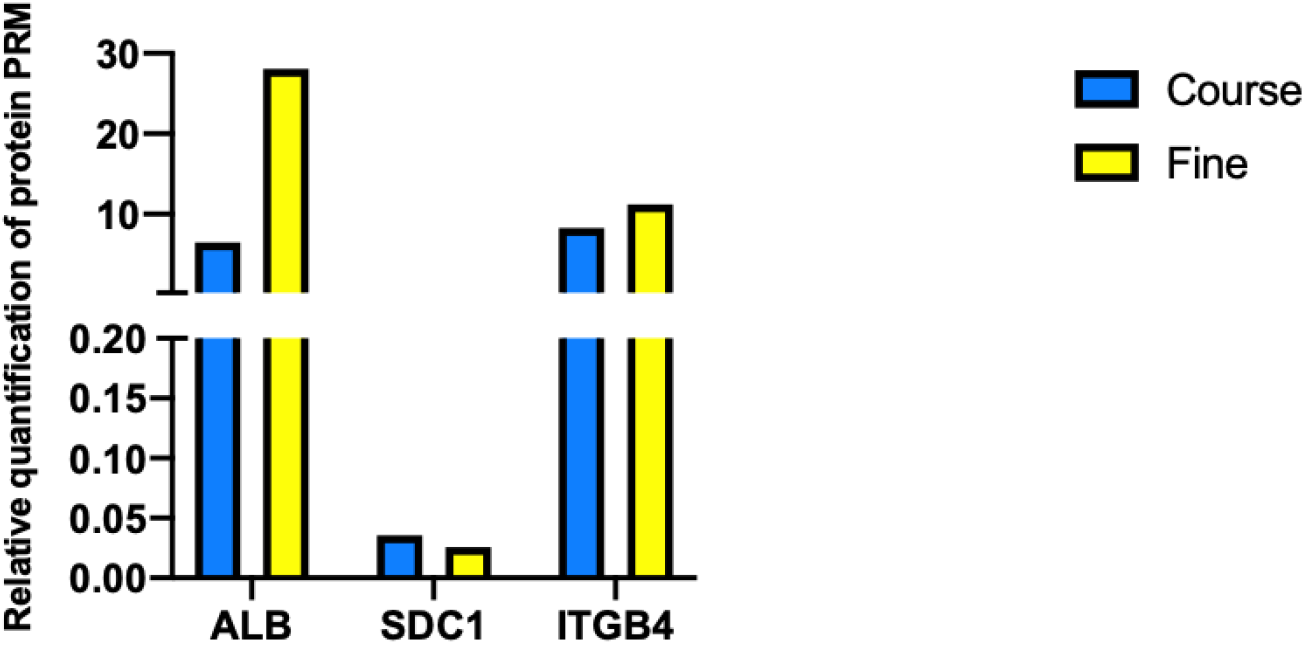
PRM validation of protein. The expression of CT_LCG and FT_LCG was verified by PRM, blue: coarse, yellow: fine.

**Table 3.** Relative quantification of differential protein PRM

| Protein Name | Gene Name | average_Coarse | average_Fine | Ratio_Coarse/Fine | Regulation |
| --- | --- | --- | --- | --- | --- |
| NP_851335.1 | ALB | 6.462822572 | 28.0845567 | 0.230120156 | up |
| XP_017911091.1 | SDC1 | 0.035479421 | 0.025566521 | 1.387729696 | down |
| XP_024835563.1 | ITGB4 | 8.26512228 | 11.22717668 | 0.736171035 | up |

### Protein- protein Interaction (PPI) networks

PPI regulatory network from the perspective of bioinformatics, which is helpful to comprehend the molecular mechanism of potential proteins involved in cashmere fineness regulation. One of the important ways for proteins to function is to interact with other proteins and exert biological regulation via protein-mediated pathways or the formation of complexes. In the PPI interaction network (Fig.5), highly aggregated proteins may exhibit the same or similar functions and perform biological functions via synergies. Protein ID: XP_005220838(*TOP2A*), NP_001069332.1(*PLA2G4A*) and XP_005227352.1(*CTSD*)have very high connectivity. *ALB*, *MGAM* and *ITGB4* are connect with *TOP2A*, *PLA2G4A* and *CTSD*. In addition, SDC1 and ALB cooperate to form PPI to regulate cashmere fineness. Therefore, they are candidates for further research.

**Fig.5.**
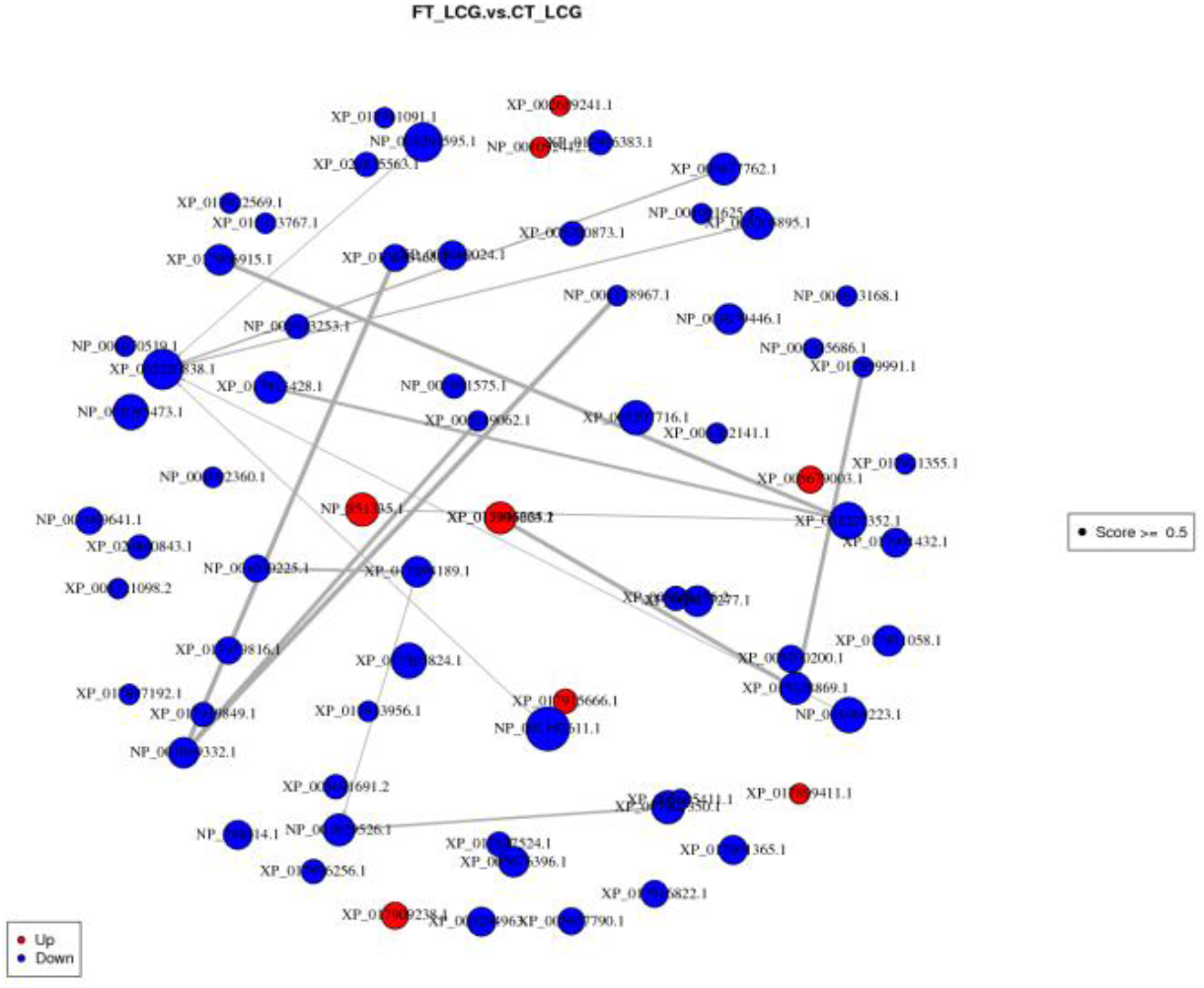
The circles represent differentially expressed proteins, and the lines correspond to protein-protein interactions. The color of the circle indicates the difference in protein expression (blue: down-regulated,red: up-regulated), and the size of the circle indicates the degree of protein connectivity (the numbers of proteins that directly interact with a particular protein).

## Conclusion

Among the 3999 identified proteins, 117 differentially expressed proteins were found. The differential expressions of *ALB*, *SDC1* and *ITGB4* in CT-LCG and FT-LCG were quantitatively identified by TMT. Then it was found that *ALB* and *SDC1* were consistent with TMT results through PRM verification. Meanwhile, *ITGB4* can be further studied as a promising protein. In the future, we will further study the regulation effect of differentially expressed protein genes on cashmere fineness. These findings will provide an important resource for further study on the regulatory functions of cashmere protein genes. *ALB*, *SDC1* and *ITGB4* genes may play a key role in cashmere fineness. This analysis provides a theoretical basis for the further study of cashmere fineness.

## Discussion

In this study, we employed a TMT-based quantitative proteomics strategy to study proteins that affect the thickness of Liaoning cashmere. Three different proteins were found between the FT_LCG and CT_LCG. We used the PRM approach to validate the differences that we found in the levels of candidate biomarker proteins, with one protein showing a similar upward or downward trend as observed by the TMT method. Currently, proteomic identification relies on comprehensive, well-annotated protein sequence databases. PRM analysis was established to validate and quantify peptides specific to selected species. The identified peptides of the selected species were verified and quantified by using PRM. Determination of protein in cashmere by PRM[20].

we identified the differential expression of 117 proteins. Among them, three proteins ALB, ITGB4 and SDC1 were significantly different, which had a certain effect on the growth of cashmere. ALB (Albumin) gene is an effective carrier tool. In Malhotra’s experiment, it was found that siRNA gene therapy with an albumin carrier, inhibit the expression of the genes involved to treat cancer[21]. In Li’s experiment, the new H11-albumin-rTTA transgenic mice were found to be a powerful and effective tool for manipulating gene expression in the liver in both spatial and temporal conditions[22]. Lorraine’s study evaluated the potential value of albuminuria as a “biomarker” for acute kidney injury (AKI) and examined whether AKI induces expression of normally silenced albumin genes in the kidney. The results showed that albuminuria had important practical value as a biomarker of acute renal tubular injury[23]. However, no relevant literature has been found on the growth of ALB gene in cashmere, which needs further study. SDC1 is a marker of epithelial-mesenchymal transformation (EMT) and can be used to evaluate tumor prognosis. SDC1 has different roles in different types of cancer[28]. In the CUI study, it was found that the overexpression frequency of SDC1 in breast cancer tissues was higher than that in normal tissues, which may be used as a potential prognostic indicator of breast cancer[29]. Yu found that SDC1 was abnormally upregulated in clinical hepatocellular carcinoma samples. Furthermore, the significance of SDC1-PI3K/ Akt signaling pathway in cisplatin resistance in HCC was emphasized[30]. The study on the growth of SDC1 protein in cashmere has not been found, which can be further studied. Members of the Integrin β (ITGB) superfamily play important roles in a variety of biological functions in a variety of cancers. ITGB4 has proven applicability in cancers of the colon, prostate, esophagus, lung, kidney, uterus, tongue, bladder and liver. Overall, ITGB4 is a reliable marker for the simultaneous detection and diagnosis of lymphatic vascular invasion (LVI) and neurological invasion (PNI). It also inhibits allergic inflammation and regulates wound repair and antioxidant capacity of airway epithelial cells [45]. ITGB4 and ITGB6 overexpression is associated with up-regulation of Notch signaling pathway, and overexpression is significantly associated with progression of AJCC stage and histological grade as well as worsening prognosis in pancreatic cancer patients[46]. ITGB4 is overexpressed in pancreatic cancer and promotes cell scattering and motility[47]. In Wilkinson’s prostate cancer study, ITGB4 gene was differentially expressed and involved in the genetic regulation of the crown of prostate cancer[48]. In Gao’s study, ITGB4 as a carrier, prostate-specific membrane antigen (PSMA) may promote angiogenesis of glioblastoma cell tumor (GBM) through its interaction with ITGB4 and activation of NF-κB signaling pathway[49]. In Li’s study, ITGB4 was found to be a new prognostic factor for colon cancer, which may become a therapeutic target and prognostic indicator for individualized treatment of colon cancer[25]. It was found in the expression and prognosis analysis of ITGA11, ITGB4 and ITGB8 in human NSCLC. ITGA11, ITGB4 and ITGB8 may be potential biomarkers and therapeutic targets[26]. ITGB4 immunotargeting is a promising therapeutic strategy for inhibiting tumor growth and reducing metastasis in RWAN’s integrin β4-targeted tumor immunotherapy[27].Studies on ITGB4 gene in sheep have only found bullosa epidermis laxity in sheep. ITGB4 is located in this region and is considered as the best candidate gene for localization and function in the combined detection of etiological mutations in sheep epidermal lytic bullous association disease by GWAS and RNA-SEQ[24]. A novel mutation of Fabre in sheep ITGB4 enhances the role of this gene in the etiology of bullosa epidermis and is an important molecular tool for improving Vendeen breed selection protocol management to limit the spread of the disease[50]. Similarly, the ITGB4 gene has not been found to be relevant in cashmere growth studies, which needs further study.

Keratin-associated proteins (KAP) are structural components in cashmere fibers, and mutations in some KAP genes are associated with many properties of cashmere fibers[31]. In an analysis of a new member of Liaoning cashmere goat keratin family (keratin 26), it was found that keratin 26 inhibited cashmere growth and was related to the regress and rest stages of hair follicles[32]. The molecular characteristics and patterns of keratin-related protein 11.1 gene expression in Liaoning cashmere goats indicate that KAP11.1 gene may regulate fiber diameter[33]. Jin’s study found that differential KAP7.1 and KAP8.2 gene expression levels in primary and secondary hair follicles may control fiber diameter in the context of wool and cashmere formation [34]. Keratin plays an essential role in cashmere growth and hair follicle development.

Moreover, we identified 173 important GO terms and 97 significant KEGG pathways. In our study, GO enrichment analysis of protein genes was mainly clustered in 20 periods(q-value < 0.05): lysosome, lipid catabolic process, carboxylic ester hydrolase activity, exopeptidase activity, plasma membrane part, cornified envelope assembly, connexin complex, omega peptidase activity, cobalamin transport, phosphohistidine phosphatase activity, peptidyl-histidine dephosphorylation transcytosis, intraciliary transport involved in cilium morphogenesis, 1-alkyl-2-acetylglycerophosphocholine esterase activity, integral component of plasma membrane, cysteine-type endopeptidase activity, membrane region, primary metabolic process, hydrolase activity, acting on ester bonds and hydrolase activity. Among them, lysosome is a highly significant pathway associated with cashmere growth. *ALB*, *SDC1* and *ITGB4* are enriched in Lysosome pathway. In the past decade, great progress has been made studying the molecular mechanisms controlling hair follicle cycle, especially the related signaling pathways[35,36]. GO and KEGG enrichment analyses of DE mRNAs and DE lncRNAs was to positively regulate typical Wnt signaling pathways and regulate protein processing and metabolism with significant enrichment[37]. In Zhao’s study showed positive regulation of many important pathways related to keratin and cell proliferation and differentiation, such as typical Wnt signaling pathways[38]. In the Bai experiment, the regulatory network of lncRNA in the Wnt signaling pathway in the secondary hair follicles of cashmere goats was studied[39]. We think that Wnt signaling pathway may be concerned with cashmere fineness.

The results showed that ALB and SDC1 were effective in the fineness growth of Liaoning cashmere through TMT combined with PRM. These differential proteins may be important for cashmere growth. However, the biological mechanisms of these protein genes need to be further investigated.

### Statement

In this work, we identified 117 differentially expressed proteins in CT_LCG (course type) and FT_LCG (fine type). To verify protein genes differentially expressed in LCG, we performed PRM validation on three candidate proteins (ALB, SDC1 and ITGB4) in CT-LCG and FT-LCG. his study sought to elucidate the molecular mechanism of differential proteins regulating cashmere fineness of Liaoning cashmere goats by using TMT quantitative proteomics analysis. Differentially expressed proteins ALB and SDC1 may regulate cashmere fineness, ITGB4 can be further studied as a promising protein. They can be used as key genes to lay a foundation for the study of cashmere fineness of Liaoning cashmere goats.

## Ethics approval and consent to participate

All Liaoning cashmere goat experiments used in this study were conducted in accordance with the guidelines of the Laboratory Animal Management Committee of Shenyang Agricultural University.

## Consent for publication

The author agrees to publish.

## Availability of data and material

The data that support the findings of this study are available from the corresponding author upon reasonable request.

## Author Contributions

Data curation, ZXB; Formal analysis, YNX and GM; Funding acquisition, ZYW; Investigation, ZXB; Methodology, ZYW; Project administration, ZYW; Resources, WDC and XJZ; Software, ZXB, YTQ, YZ, RC, YGS and YZW; Supervision, ZYW; Validation, ZXB; Visualization, ZYW; Writing – original draft, ZXB; Writing – review & editing, ZXB and ZY

## Acknowledgements

1. Thanks to the teachers and students of Shenyang Agricultural University in the technical help, including instruments and equipment and their related experimental materials, collaborative experimental work, to provide useful inspiration, suggestions, guidance, review, undertake some auxiliary work, etc.
2. Thanks to the Foundation for helping the Grants from the National Natural Science Foundation of China(NO.31802038), the 69th batch of China Postdoctoral Science Foundation (Regional Special Support Program) (NO. 2021M693859), Youth seedling project of Liaoning Provincial Department of Education, China(LSNQN201905) and Liaoning Provincial Department of science and technology, agricultural key issues and industrialization project (2020JH2/10200029).

